# Improving Small Molecule pK_*a*_ Prediction Using Transfer Learning with Graph Neural Networks

**DOI:** 10.1101/2022.01.20.476787

**Authors:** Fritz Mayr, Marcus Wieder, Oliver Wieder, Thierry Langer

**Affiliations:** Department of Pharmaceutical Sciences, Pharmaceutical Chemistry Division, University of Vienna, Althanstrasse 14, 1090 Vienna, Austria

## Abstract

Enumerating protonation states and calculating micro-state pK_*a*_ values of small molecules is an important yet challenging task for lead optimization and molecular modeling. Commercial and non-commercial solutions have notable limitations such as restrictive and expensive licenses, high CPU/GPU hour requirements, or the need for expert knowledge to set up and use. We present a graph neural network model that is trained on 714,906 calculated mico-state pK_*a*_ predictions from molecules obtained from the ChEMBL database. The model is fine-tuned on a set of 5,994 experimental pK_*a*_ values significantly improving its performance on two challenging test sets. Combining the graph neural network model with `Dimorphite-DL`, an open-source program for enumerating ionization states, we have developed the open-source Python package pkasolver, which is able to generate and enumerate protonation states and calculate micro-state pK_*a*_ values with high accuracy.

## Introduction

The acid dissociation constant (K_*a*_), most often written as its negative logarithm (pK_*a*_), plays a major role in the field of molecular modeling, as it influences the charge, tautomer configuration, and overall 3D structure of molecules with accessible protonation states in the physiological pH range. All these factors further shape the mobility, permeability, stability, and mode of action of substances in the body [1]. In case of insufficient or missing empirical data, the correct determination of pK_*a*_ values is thus essential to correctly predict the aforementioned molecular properties.

Authors and studies disagree on the exact percentage of drugs with ionizable groups, but a conservative estimate suggests that at least two-thirds of all drugs contain one or more ionization groups (in a pH range of 2 to 12) [2]. The importance of pK_*a*_ predictions for drug discovery has been widely recognized and has been the topic of multiple blind predictive challenges (most notable the Statistical Assessment of Modeling of Proteins and Ligands (SAMPL) series SAMPL6 [3], SAMPL7, [4] and ongoing SAMPL8 ^1^ challenge.

Multiple methods have been developed to estimate pK_*a*_ values of small molecules, ranging from physical models based on quantum chemistry calculations [5, 6] and/or free energy calculations [7, 8] to empirical models based on linear free energy relationships using the Hammet-Taft equation or more data driven methods using quantitative structure-property relationship (QSPR) and machine learning (ML) approaches like deep neural network or random forest models [4, 9–11]. In general empirical methods require significantly less computational effort than their physics-based counterparts once they are parameterized but require a relatively large number of high-quality data points as training set [4].

In recent years, machine learning methods have been widely applied to predict different molecular properties including pK_*a*_ predictions. Many of these approaches learn pK_*a*_ values on fingerprint representations of molecules [10, 12]. The pK_*a*_ value of an acid and conjugate base pair is determined by the molecular structure and the molecular effects on the reaction center exerted by its neighborhood, including mesomeric, inductive, steric, and entropic effects [13]. Ideally, these effects should be included and encoded in a suitable fingerprint or set of descriptors. For many applications, extended-connectivity fingerprints (ECFPs) in combination with molecular features have proven to be a suitable and powerful tool to learn structure-property relationships [14, 15].

The emergence of graph neural networks (GNNs) has shifted some focus from descriptors and fingerprints designed by domain experts to these emerging deep learning methods. GNNs are a class of deep learning methods designed to perform inference on data described by graphs and provide straightforward ways to perform node-level, edge-level, and graph-level prediction tasks [16–18]. GNNs are capable of learning representations and features for a specific task in an automated way eliminating the need for excessive feature engineering ([19]). Another aspect of their attractiveness for molecular property prediction is the ease with which a molecule can be described as an undirected graph, transforming atoms to nodes and bonds to edges encoded both atom and bond properties. GNNs have proven to be useful and powerful tools in the machine learning molecular modeling toolbox [19, 20].

Pan et al. [21] have shown that GNNs can be successfully applied to pK_*a*_ predictions of chemical groups of a molecule, outperforming more traditional machine learning models relying on human-engineered descriptors and fingerprints, developing `MolGpka`, a web server for predicting pK_*a*_ values. `MolGpka` was trained on molecules extracted from the ChEMBL database [22] containing predicted pK_*a*_ values (predicted with ACD/Labs Physchem software^2^). Only the most acidic and most basic pK_*a*_ values were considered for the training of the GNN models.

The goal of this work was to extend the scope of predicting pK_*a*_ values for independently ionizable atoms (realized in `MolGpka`) and develop a workflow that is able to enumerate protonation states and calculate the corresponding pK_*a*_ values connecting them (sometimes referred to as ‘sequential pK_*a*_ prediction‘). To achieve this we implemented and trained a GNN model that is able to predict values for both acidic and basic groups by considering the protonated and deprotonated species involved in the corresponding acid-base reaction. We trained the model in two stages. First, we started by pre-training the model on calculated microstate pK_*a*_ values for a large set of molecules obtained from the ChEMBL database [22]. The pre-trained model already performs well on the two independent test sets used to measure the performance of the trained models. To improve its performance we fine-tuned the model on a small training set of molecules for which experimental pK_*a*_ values were available. The fine-tuned model shows excellent and improved performance on the two test sets.

We have implemented the training routine and prediction pipeline in an open-source `Python` package named `pkasolver`, which is freely available and can be obtained as described in the **Code and data availability** section. We also provide a ready-to-use `Google Colab Jupyter` notebook which includes the trained models and can be used to predict pk_*a*_ values for molecules without locally installing the package (for further information see the **Code and data availability** section) [23].

## Results & Discussion

We will start by discussing the performance of the model on the validation set of the ChEMBL data set (which contains micro-state pK_*a*_ values calculated with `Epik` on a subset of the ChEMBL database) and the two independent test sets: the Novartis test set (280 molecules) and the Literature test set (123 molecules). This will be followed by a discussion of the fine-tuned model on its validation set, on both test sets, and on the ChEMBL data set. Finally, we will discuss the developed `pkasolver` package and its use cases.

Performance of the different predictive models is subsequently reported using the mean absolute error (MAE) and root mean squared error (RMSE). For each metric (MAE and RMSE) the median value from 50 repetitions with different training/validation set splits is reported and the 90% confidence interval is shown. To visualize training results a single training run was randomly selected and the results on the validation set plotted.

### Pre-training model performance

The initial training of the GNN model was performed using the ChEMBL data set (microstate pK_*a*_ values calculated with `Epik`). Figure S.I.3 **A** shows the results of the best performing model on the hold-out validation set. The MAE and RMSE are 0.29 [90% CI: 0.28; 0.31] and 0.45 [90% CI:0.44;0.49] pK_*a*_ units showing an excellent accuracy across the reference pK_*a*_ values. The kernel density estimates (KDE) of the distribution of the reference and predicted pK_*a*_ values shown in Figure S.I.3 **A** highlights the ability of the GNN to correctly predict pK_*a*_ values throughout the investigated pH range.

The performance of the trained GNN model was assessed on two independent test sets: the Novartis and the Literature test set (both test sets are described in detail in the **Methods** section) [10]. The trained model performs well on both test sets with a MAE of 0.62 [90% CI:0.57;0.67] and a RMSE of 0.97 [90% CI:0.89;1.10] pk_*a*_ units on the Literature test set and a MAE of 0.82 [90% CI:0.77;0.85] and a RMSE of 1.13 [90% CI:1.05;1.21] pk_*a*_ units on the Novartis test set (shown in Figure S.I.2). The performance is comparable to the performance of `Epik` and `Marvin` on both test sets (shown in Table 1).

**Table 1.** Performance of state-of-the-art knowledge-based approaches and commercial software solutions to predict pK_*a*_ values on the Novartis and Literature test sets are shown. For each data set, the mean absolute error (MAE) and root mean squared error (RMSE) is calculated. For `MolGpka`, `Epik`, and `pkasolver` the median value and the 90% confidence interval are reported. ^1^ used a random forest implementation with 1,000 estimators and the FCFP6 fingerprint. Values for the best performing random forest implementation are shown. ^2^ `Epik` identified different protonation centers than were reported in the data sets for the Novartis data set for 26 out of 280 molecules. These molecules were excluded from the MAE and RMSE calculation for `Epik`. ^3^ values were obtained from Baltruschat and Czodrowski [10]. ^4^ the reason for the large confidence interval is the incorrect prediction for a single molecule (Isomeric Smiles: CCNC) by `MolGpka` with an error of 8.86 pK_*a*_ units.

| Model | Novartis data set |  | Literature data set |  |
| --- | --- | --- | --- | --- |
|  | MAE | RMSE | MAE | RMSE |
| Random Forest<sup>1,3</sup> | 1.15 | 1.51 | 0.53 | 0.76 |
| ChemAxon Marvin (V20.1.0)<sup>3</sup> | 0.86 | 1.17 | 0.57 | 0.87 |
| MolGpKa [21] | 0.87 [0.77;0.97] | 1.27 [1.08;1.45] | 0.49 [0.40;0.65] | 1.00 [0.56;1.53]<sup>4</sup> |
| Epik<sup>2</sup> [21] | 0.83 [0.75;0.91] | 1.16 [1.06;1.26] | 0.58 [0.48;0.67] | 0.92 [0.74;1.12] |
| pkasolver | 0.71 [0.64;0.74] | 0.93 [0.85;0.97] | 0.52 [0.49;0.56] | 0.82 [0.76;0.86] |

### Fine-tuned model performance

While the performance on the test sets of the pre-trained model was already acceptable we were able to further increase model accuracy by fine-tuning the pre-trained model using a data set of experimentally measured pK_*a*_ values. The performance of the fine-tuned model on the validation set of the experimental data set is shown in S.I.3 **B**. The median performance of the fine-tuned model was improved from a RMSE of 0.97 [90% CI:0.89;1.10] to 0.82 [90% CI:0.76;0.88] pK_*a*_ units on the Literature test set and from a RMSE of 1.13 [90% CI:1.05;1.21] to 0.93 [90% CI:0.85;0.97] pK_*a*_ units on the Novartis test set (shown in Figure 1).

**Figure 1.**
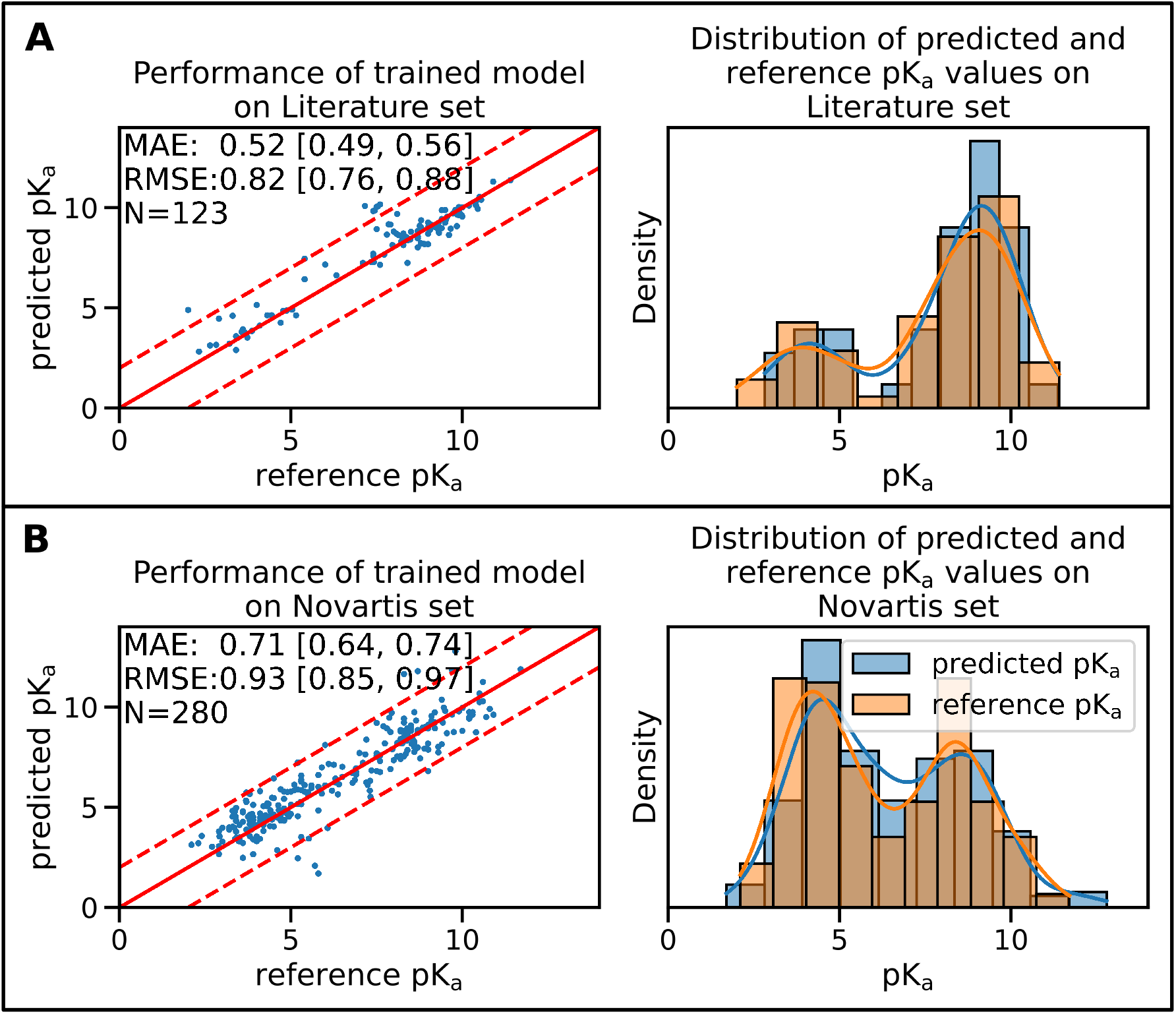
The fine-tuned GNN model is able to predict the pK_*a*_ values of the Novartis and Literature test set with high accuracy. Panel **A** shows the performance of the fine-tuned model (initially trained with the ChEMBL data set and subsequently fine-tuned on the experimental data set) on the Literature test set. Panel **B** shows the performance of the same model on the Novartis test set. The solid red line in the scatter plot indicates the ideal behavior of the reference and predicted pK_*a*_ values, the dashed lines mark the ±1 pk_*a*_ unit interval. Mean absolute error (MAE) and root mean squared error (RMSE) is shown, the values in bracket indicate the 90% confidence interval calculated from 50 repetitions with random training/validation splits. ***N*** indicates the number of investigated samples.

In order to avoid model performance degradation on the ChEMBL data set we randomly added molecules from the ChEMBL data set during the fine-tuning workflow. Adding molecules from the ChEMBL data set to restrict model parameters and avoid overfitting decreased the performance of the fine-tuned model on the ChEMBL data set only slightly (shown in Figure S.I.4). This was necessary since previous attempts without regularization showed decreased accuracy of the fine-tuned model in regions outside the limited pH range of the experimental data set while improving the performance on the test sets (details to the pH range of both the ChEMBL and experimental data set are shown in Figure S.I.5). An example of the performance of the fine-tuned model on the ChEMBL data set without regularization is shown in Figure S.I.6.

To set the performance of the fine-tuned model in context we compare its performance with two recent publications investigating pK_*a*_ predictions using machine learning. In Table 1 the results are summarized for the methods presented in both Baltruschat and Czodrowski [10] and Pan et al. [21]. We extracted data from these publications where appropriate and recalculated values where needed. Pan et al. [21] split the reported results into basic and acidic groups making it necessary to recalculate the values reported there for `MolGpka` and `Epik`, the values for Marvin were taken directly from reference [10] (reported values were calculated without confidence interval). The fine-tuned GNN model (called `pkasolver` in Table 1) performs on a par with the best performing methods reported there.

It is difficult to rationalize MAE/RMSE differences between different methods/models shown in Table 1) since training sets and methods are different. The small difference in performance between `pkasolver` and `MolGpka` could be attributed to the transfer learning routine which added experimentally measured pK_*a*_ values. The random forest model was trained on significantly less data (only on the 5,994 pk_*a*_ values present in the experimental data set) than either `pkasolver` or `MolGpka` yet performs comparably to both on the Literature data set while significantly worse on the Novartis data set. This might highlight the complexity of the Novartis data set, an observation previously made and investigated in Pan et al. [21].

Both `Epik` and `Marvin` perform well on both test data sets. It is surprising that `pkasolver` can slightly outperform `Epik`, even though its initial training was based on data calculated by `Epik`. We think this emphasizes the potential of transfer learning as used in this work and data-driven deep learning in general.

### Sequential pK_*a*_ predictions with pkasolver

Combining the trained GNN model with Dimorphite-DL, a tool that identifies potential protonation sites, enabled us to perform sequential pK_*a*_ predictions. A detailed description of this approach is given in the **Detailed methods** section. We investigated multiple mono and polyprotic molecules for qualitative and quantitative (in reasonable bounds) agreement between prediction and experimental data. The results for the investigate systems were of excellent consistency. The list of molecules that we tested is included in the `pkasolver` repository.

We want to discuss the results generated by `pkasolver` on a challenging molecule: ethylenediaminetetraacetic acid, better known as EDTA. The protonation states and molecules are shown in Figure S.I.7. The correct protonation states are identified and the pK_*a*_ values for the 6 protonation states estimated are 1.49, 1.93, 2.28, 2.57, 6.05, and 9.44. Comparing this with the experimentally determined pK_*a*_ values of 0.0, 1.5, 2, 2.66, 6.16, and 10.24 the values are in good relative agreement [24]. Only the first and last pK_*a*_ estimates are noticeable too high and too low respectively. This highlights a limitation of `pkasolver`. The range of pK_*a*_ values present in the training data was between zero and 14, and predictions of pK_*a*_ values will stay in this range.

## Conclusion

We have shown that GNNs can be used to predict mico-state pK_*a*_ values and achieve excellent performance on two external test sets. Training the GNN model in two stages with a pre-training phase using a large set of molecules with calculated pK_*a*_ values and a fine-tuning phase on a small set of molecules with experimentally measured pK_*a*_ values improves the performance of the GNN model significantly. This performance boost is especially noteworthy on the challenging Novartis test set (the RMSE was decreased from 1.18 [1.05;1.27] to 0.93 [0.85;0.97] pk_*a*_ units). A direct comparison with other software solutions and machine learning models on the two test sets shows that the fine-tuned GNN model performs consistently on a par with the best results of other commercial and non-commercial tools.

We have implemented `pkasolver` as an open-source and free-to-use `Python` package under a permissive licence (MIT licence). We provide an easy-to-use and accurate tool that enumerates protonation states and calculates micro-state pK_*a*_ values closing a gap in the computational chemistry toolbox. By providing the original training data, the fine-tuned model parameters, and training routines we encourage the community to build on this work.

## Code and data availability

- Python package used in this work (release v0.2) and Colabs Jupyter notebook link: https://github.com/mayrf/pkasolver
- Data and notebooks to reproduce the plots/figures (release v0.1): https://github.com/wiederm/pkasolver-data

## Author Contributions

Conceptualization: FM, OW, TL, and MW; Methodology: FM, OW, and MW; Software: FM and MW; Investigation: FM and MW; Writing–Original Draft: FM, OW, and MW; Writing–Review&Editing: FM, OW, TL, and MW; Funding Acquisition: MW, TL; Resources: TL; Supervision: MW, TL.

## Acknowledgments

MW is grateful for discussions with David Bushiri, John Chodera, Josh Fass, Nils Krieger, Magdalena Wiercioch, Steffen Hirte, Thomas Seidel, and the Tautomer Consortium, specifically Paul Czodrowski, Brian Radak, Woody Sherman, David Mobley, Christopher Bayly, and Stefan Kast. MW, OW, FM, and TL are grateful for the help of Gerhard F. Ecker and his group members who performed the reference pK_*a*_ calculations with `Epik` for the ChEMBL data set.

## Detailed methods

### Data set generation and pre-processing

Four different data sets were used in this work: the ChEMBL data set, the experimental data set, the Novartis data set and the Literature data set.

The ChEMBL data set used for pre-training was obtained from the ChEMBL database using the number of Rule-of-Five violations (set to a maximum of one violation) as filter criteria [22, 25]. For each of the molecules, a pK_*a*_ scan for the pH range between zero and 14 was performed using the Schrodinger tool `Epik` [26, 27] (Version-2121-1). The sequential pK_*a*_ scan indicated for 320,800 molecules one or multiple protonation state/s, resulting in a total of 729,375 pK_*a*_ values. For each pK_*a*_ value, `Epik` further indicated the protonation center using the atom index of the heavy atom at which either a hydrogen is attached or removed.

To perform transfer learning we obtained a second data set with experimental pK_*a*_ values. This data set (subsequently called ‘experimental data set‘) was developed by Baltruschat and Czodrowski [28] and can be acquired from their GitHub repository^3^. For a detailed description of the curating steps taken to generate this data set, we point the reader to the **Methods** section of [28]. The experimental data set consists of 5,994 unique molecules, each with a single pK_*a*_ value and an atom index indicating the reaction center. Some of the molecules had to be corrected to obtain their protonation state at pH 7.4 (examples shown in S.I.1).

To test the performance of the models, two independent data sets were used, which were provided and curated by Baltruschat and Czodrowski [28]. The Literature data set contains 123 compounds collected by manual curating the literature. The Novartis data set contains 280 molecules provided by Novartis [29]. For each molecule, a pK_*a*_ value and atom index indicating the reaction center was provided. To avoid training the model on molecules present in the Literature or Novartis data set we filtered the ChEMBL data set using the InChIKey and canonical SMILES strings of the neutralized molecules as matching criteria. 50 molecules were identified and removed from the ChEMBL data set. All checks were performed using `RDKit` [30].

### Enumerate protonation states during training/testing

The goal of calculating microstate pK_*a*_ values is to find the pH value at which the concentration of two molecular species is equal. To do this efficiently, we provide as input the protonated and deprotonated molecular species of the acid-base pair for which we want to calculate the pK_*a*_ value (the Brønsted acid/base definitions are used here and subsequently [31]). This approach enables a consistent treatment of acids and bases with a single data structure (the acid-base pair).

This workflow made it necessary that we generate the molecular species at each protonation state starting from the molecule at pH 7.4 by removing or adding hydrogen to the reaction center (which was calculated by `Marvin` for the experimental, Novartis, and Literature data set and `Epik` for the ChEMBL data set). We do this by sequentially adding hydrogen atoms from highest to lowest pK_*a*_ for acids (i.e. at pH=0 all possible protonation sites are protonated) and removing hydrogen atoms from lowest to highest pK_*a*_ value for bases on the structure present at pH 7.4 (at pH=14 all possible protonation sites are deprotonated).

This approach presented challenges for the ChEMBL data set for which sequential pK_*a*_ values and reaction centers were calculated with `Epik`. `Epik` calculates the micro-state pK_*a*_ value on the most probable tautomeric/mesomeric structure. This leads to potential protonation states that require changes in the double bond pattern and redistribution of hydrogen. Since we do not consider tautomeric changes to the molecular structure in the present implementation such tautomeric changes can introduce invalid molecules in either the sequential removal or addition of hydrogen atoms. Whenever such molecular structures were encountered we removed these protonation states from further consideration. Additionally, we used RDKit’s sanitize function to identify cases for which protonation state changes introduce invalid atom valences. In other cases in which the protonation state change on a mesomeric structure introduces valid yet improbable molecular structures (e.g. protonating the oxygen in an amide instead of the nitrogen) we keep these structures. This reduced the number of molecules and protonation states in the ChEMBL dataset to 286,816 molecules and 714,906 protonation states. The distribution of pK_*a*_ values for the ChEMBL and experimental data set is shown in Figure S.I.5.

### Training and testing with PyTorch geometric

We use `PyTorch` and `PyTorch geometric` (subsequently abbreviated as `PyG`) for model training, testing, and prediction of pK_*a*_ values on the graph data structures [33, 34].

#### Graph data structure

A graph ***G*** is defined as a set of nodes ***V*** and edges ***E*** connecting the nodes. Each node ***v*** ∈ ***V*** has a feature vector *x*_*v*_, which encodes atom properties like element, charge, number of hydrogen, etc. as well as the presence of particular SMARTS patterns as a one-hot-encoding bit vector (all atom properties are shown in Table S.I.1). The adjacency matrix *A* defines the connectivity of the graph. *A* is defined as a quadratic matrix with ***A***_*uv*_ = 1 if there is an edge between node *u* and *v* and ***A***_*uv*_ = 0 if there is no edge between node *u* and *v*.

We used `RDKit` to generate a graph representation of the molecule with atoms represented as nodes and bonds as edges (in coordinate list format ^4^ to efficiently represent the spares matrix).

#### Graph neural network architecture

To predict a single pk_*a*_ value the graph neural network (GNN) architecture takes as input two graphs representing the conjugated acid-base pair as shown in Figure 2. Figure 2 **B** shows the high-level architecture of the used GNN.

**Figure 2.**
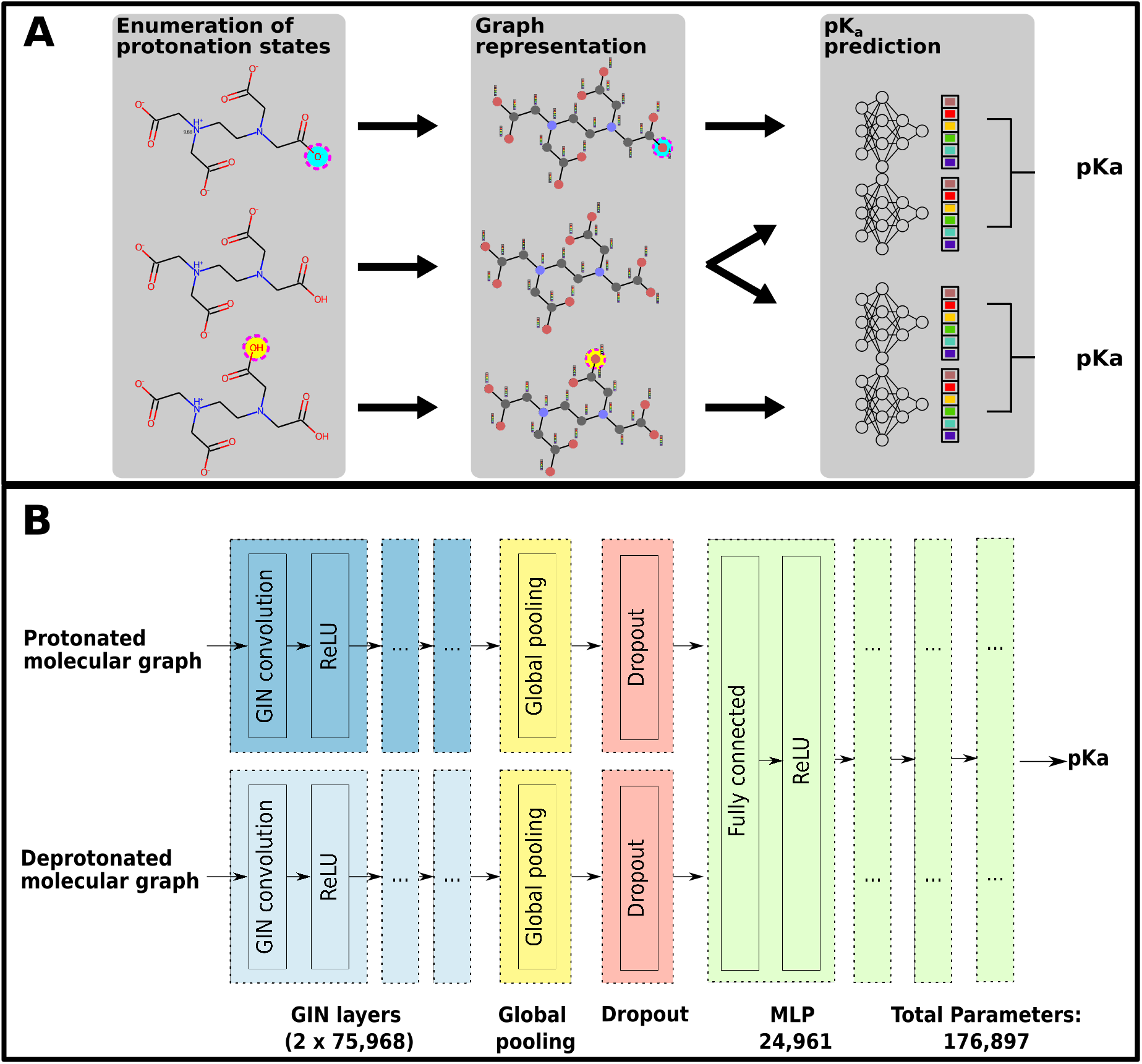
Panel A shows the general workflow used to train the GNN on microstate pK_*a*_ values for a single molecule. During the training and testing phase, each molecule was provided in the structure dominant at pH 7.4 with atom indices indicating the protonation sites and corresponding pK_*a*_ values connecting them. In the **Enumeration of protonation states** phase we generate the protonation state for each pK_*a*_ value. The molecular species for each of the protonation states are then translated in their graph representation using nodes for atoms and edges for bonds, with node feature vectors encoding atom properties in the **Graph representation** phase. In the **pK**_*a*_ **prediction** phase graphs of two neighboring protonation states are combined and used as input for the GNN model to predict the pK_*a*_ value for the acid-base pair (using the Brønsted–Lowry acid/base definition [31]). The architecture of the GNN model is shown in detail in panel **B**. For a pair of neighboring protonation states two independent GIN (graph isomorphism network) convolution layers and ReLU activation functions are used for the protonated and the deprotonated molecular graph to pass information of neighboring atoms and achieve the embedding of the chemical environment of each atom [32]. The output of the convolutional layer is summarized using a global average pooling layer, generating the condensed input for the multilayer perceptron (MLP). To add regularization and to prevent co-adaptation of neurons a dropout layer was added.

There are three phases to predict a pK_*a*_ value from a pair of molecular graphs. The first stage involves recurrently updating the node states using GIN (graph isomorphism network) convolution layers and ReLU activation functions [32]. We used 3 GIN layers with an embedding size of 64 bits each to propagate information throughout the graph and update each node with information about the extended environment. In the second stage, a global average pooling is performed to produce the embedding of the protonated and de-protonated graph, resulting in two 32 bit vectors. Concatenating the two 32 bit vectors produces the input for the third stage, the multilayer perceptron (MLP) with 3 fully connected layers (each with an embedding size of 64). To add regularization and to prevent co-adaptation of neurons a dropout layer randomly zeros out elements of the pooling output vector with *p* = 0.5 during training. Additionally, batch normalization is applied as described in [35].

#### GNN model training

Before each training run the ChEMBL and experimental data set were shuffled and randomly split in training (90% of the data) and validation set (10% of the data). To ensure that we can reproduce these splits the seed for each split was recorded.

The mean squared error (MSE) of predicted and reference pK_*a*_ values on the training data set was calculated and parameter optimization was performed using the Adam optimizer with decoupled weight decay regularization [36] as implemented in `PyTorch`.

Model performance was evaluated on the validation set and the model with the best performance was selected either for fine-tuning or further evaluation on the test data sets. The performance on the evaluation data set was calculated after every fifth epoch and the corresponding weights were saved. The learning rate for all training runs was dynamically reduced by a factor of 0.5 if the validation set performance did not change within 150 epochs (validation set performance threshold was set to 0.1).

Pre-training of the GNN was performed on the ChEMBL data set with a learning rate of 1*x*10^−3^ and a batch size of 512 molecules for 1,000 epochs. Fine-tuning was performed using the experimental data set with a learning rate of 1*x*10^−3^ and a batch size of 64 molecules for 1,000 epochs. All parameters of the GNN models were optimized during fine-tuning. To avoid overfitting to the experimental data set we added to each batch of the fine-tuning data set a randomly selected batch (1024 molecules) of the pre-training data set in the gradient calculation phase.

To calculate the confidence intervals of the model performance, pre-training and fine-tuning were repeated 50 times, each with a random training-validation set split resulting in 50 independently fine-tuned models.

### Sequential pK_*a*_ value prediction with pkasolver

We use `Dimorphite-DL` to identify the proposed structure at pH 7.0 and all de-/protonation sites for a given molecule [37].

Then we iteratively protonate each of the proposed de-/protonation sites generating a molecular pair consisting of the protonated and deprotonated molecular species (in the first iteration the deprotonated molecule is the molecule at pH 7.0). For each of the protonate/deprotonated pairs a pK_*a*_ value is calculated. The protonated structure with the highest pK_*a*_ value (but below pH 7.0) is kept and the protonation site is removed from the list of possible protonation sites. This is repeated until either (1) all protonation sites are protonated, (2) no more valid molecules can be generated, or (3) the calculated pK_*a*_ values are outside the allowed pK_*a*_ range.

To enumerate all deprotonated structures we start again with the structure at pH 7.0 and start to iteratively deprotonate each of the proposed de-/protonation sites. Here, we always keep the deprotonated structure with the lowest pK_*a*_ value that is above 7.0.

The pK_*a*_ values are calculated using 25 of the 50 fine-tuned GNN models. For each protonation state, the average pK_*a*_ value is calculated and the standard deviation is shown to enable the user to identify molecules or protonation states for which the GNN model estimates are uncertain.

We provide a ready to use implementation of `pkasolver` to predict sequential pK_*a*_ values in our GitHub repository (for further information see the **Code and data availability** section).

#### Supplementary Information

**Table S.I.1.**
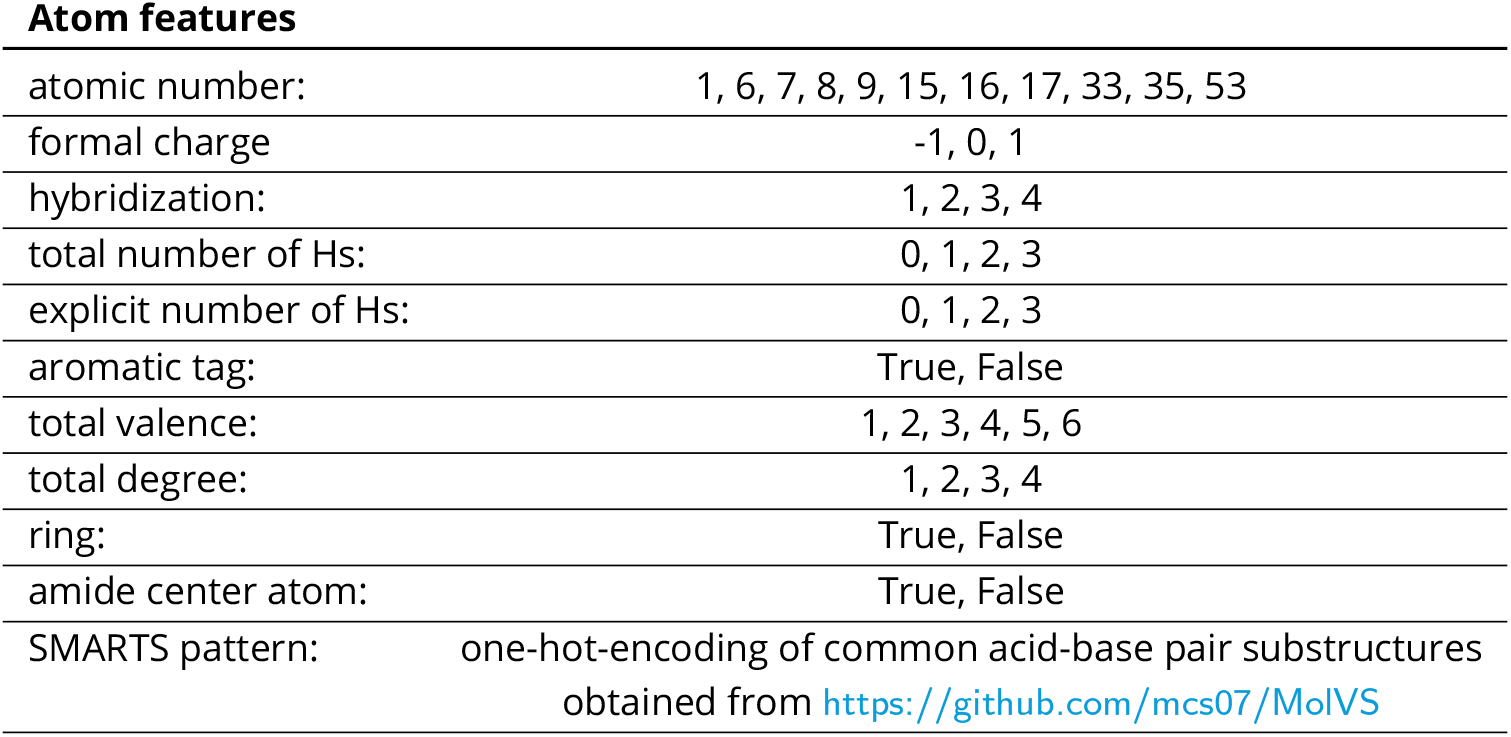
List of one-hot-encoding of atom features used for the node feature vector deposited in the node feature matrix ***X***.

**Figure S.I.1.**
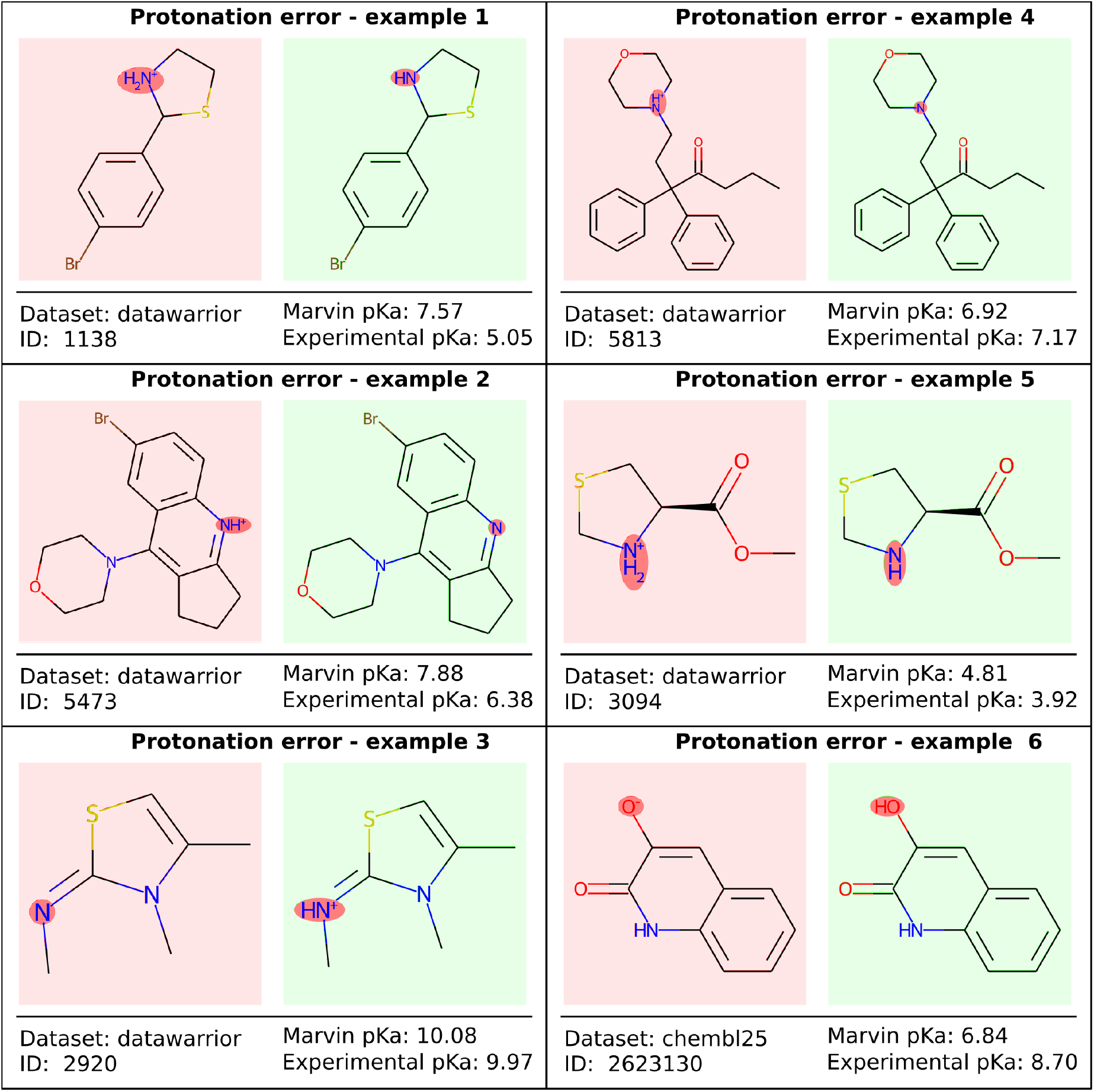
Protonation state errors in the experimental data set. This exemplary selection shows molecules from the experimental data set provided by Baltruschat and Czodrowski [10] for which the protonation state provided does not correspond to the state at pH 7.4. For examples 1, 2, 4 and 5 with experimental *pK*_*a*_ values below 7.4 protonation at the reaction center would result in highly unlikely pentavalent nitrogen. For example 3 and 6 with *pK*_*a*_ values above 7.4 deprotonation at the reaction site can not be performed because of the lack of a suitable hydrogen. These error were corrected during our data preparation.

**Figure S.I.2.**
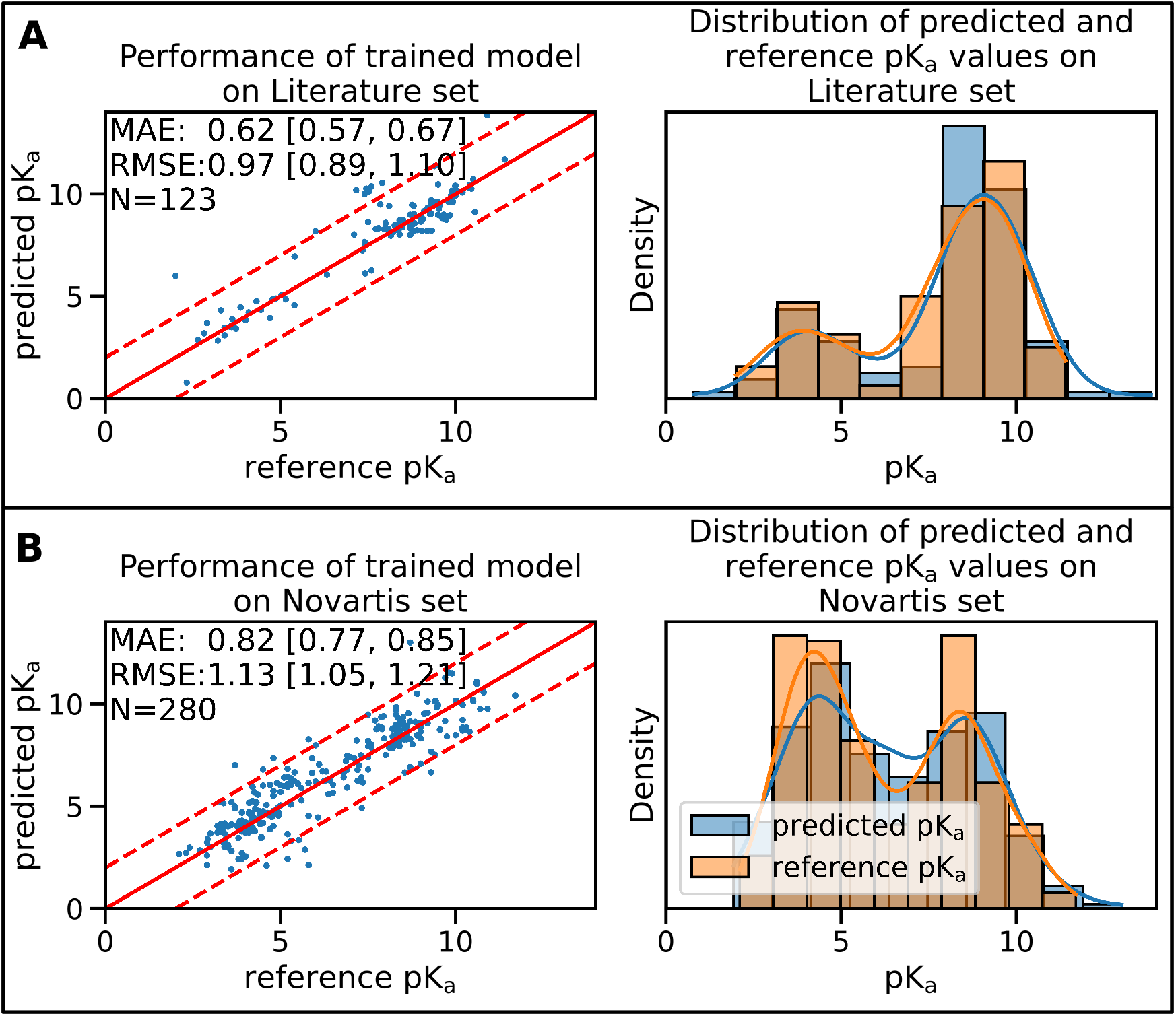
Performance of the pre-trained GNN model on the Novartis and Literature test set is shown. 50 training runs with different training/validation splits were performed and for each training run the best model was selected based on its performance on the validation set (shown here is a single, randomly selected training run). Panel **A** shows the performance of the GNN model on the Literature data set. Panel **B** shows the performance of the GNN model on the Novartis data set. The solid red line in the scatter plot indicates the ideal behavior of the reference and predicted pK_*a*_ values, the dashed lines mark the ±1 pk_*a*_ unit interval. Mean absolute error (MAE) and root mean squared error (RMSE) are shown, the values in bracket indicate the 90% confidence interval calculated from 50 repetitions with random training/validation splits. *N* indicates the number of investigated samples.

**Figure S.I.3.**
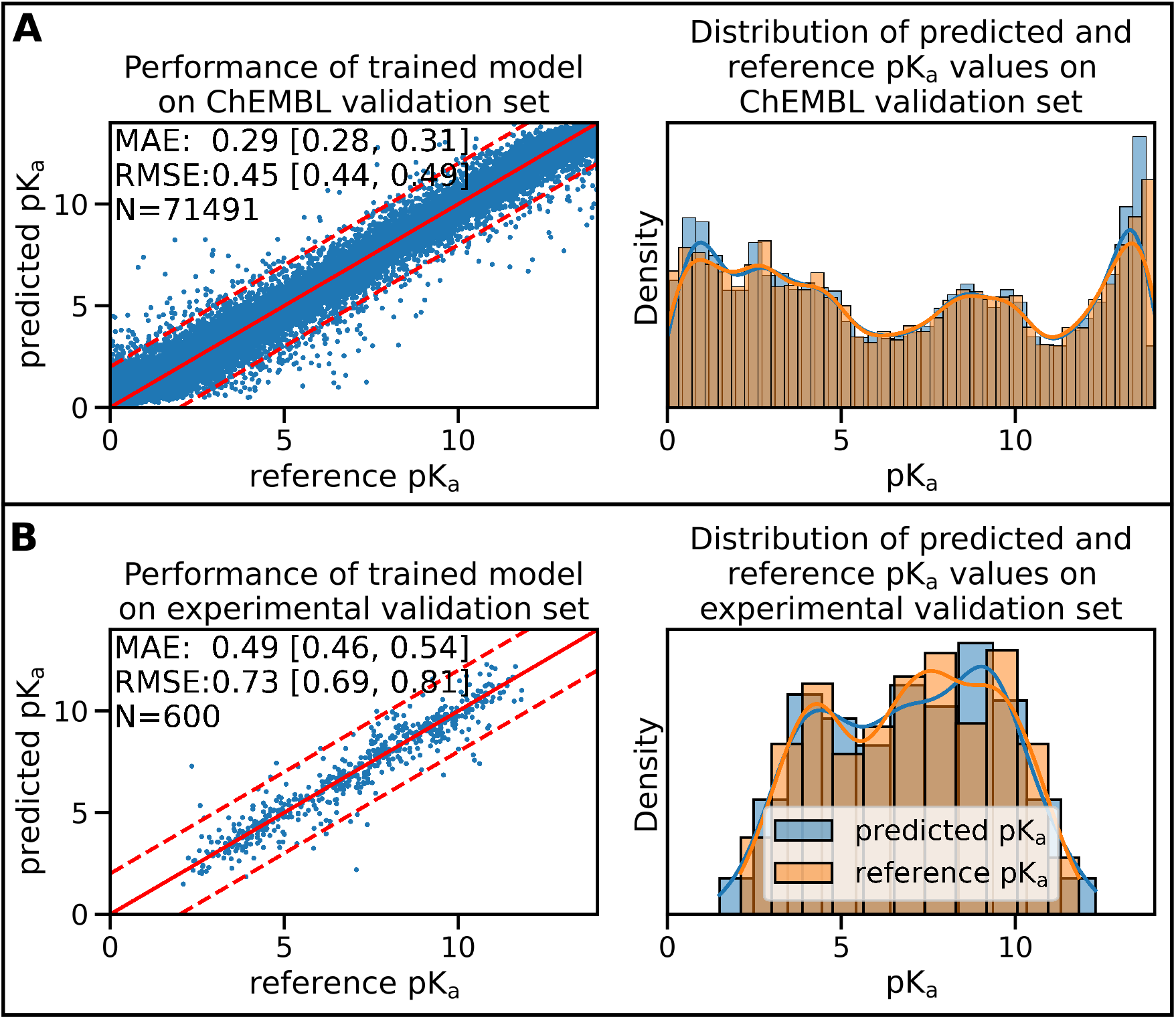
Performance of the pre-trained and fine-tuned models are shown on the respective validation sets. 50 training runs with different training/validation splits were performed and for each training run the best model was selected based on its performance on the validation set (shown here is a single, randomly selected training run). Panel **A** shows the validation set performance of the best GNN model trained on the ChEMBL data set. Panel **B** shows the validation set performance starting from the same pre-trained model after fine-tuning on the experimental training set. The solid red line in the scatter plot indicates the ideal behavior of the reference and predicted pK_*a*_ values, the dashed lines mark the ±1 pk_*a*_ unit interval. Mean absolute error (MAE) and root mean squared error (RMSE) are shown, the values in bracket indicate the 90% confidence interval calculated from 50 repetitions with random training/validation splits. *N* indicates the number of investigated samples.

**Figure S.I.4.**
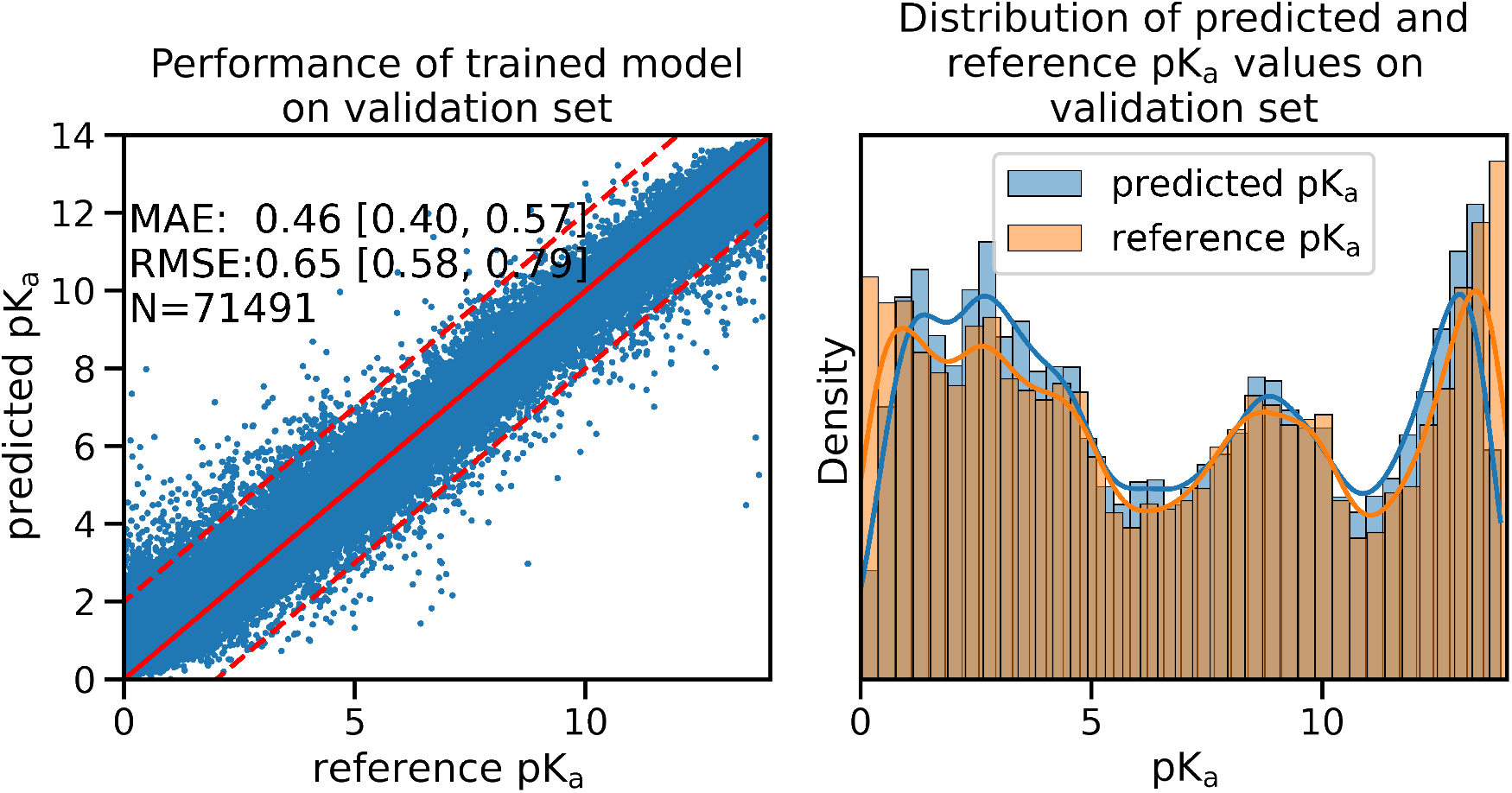
The accuracy of the fine-tuned GNN model only decreases slightly when molecules from the ChEMBL data set are used for regularization. 50 fine-tuning runs with different training/validation splits were performed, each initialized using the parameters of 50 pre-training runs, and for each training run the best model was selected based on its performance on the validation set. In order to generate a single plot we selected randomly a single fine-tuning run and generated the scatter plot with the best performing model on the validation set. The solid red line in the scatter plot indicates the ideal behavior of the reference and predicted pK_*a*_ values, the dashed lines mark the ±1 pk_*a*_ unit interval. Mean absolute error (MAE) and root mean squared error (RMSE) are shown, the values in bracket indicate the 90% confidence interval calculated from 50 repetitions with random training/validation splits. ***N*** indicates the number of investigated samples.

**Figure S.I.5.**
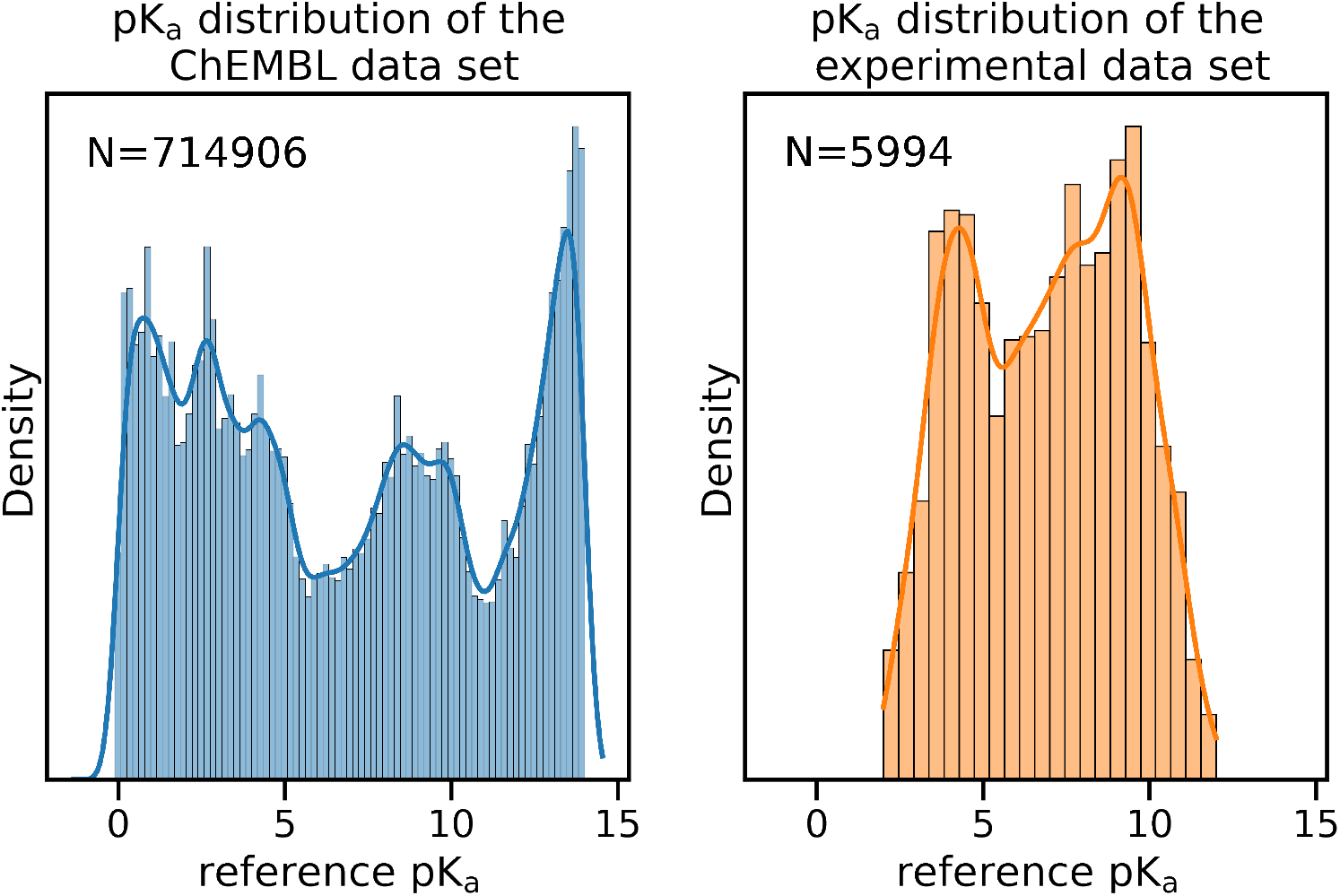
The pk_*a*_ distribution of ChEMBL and experimental data set are shown.

**Figure S.I.6.**
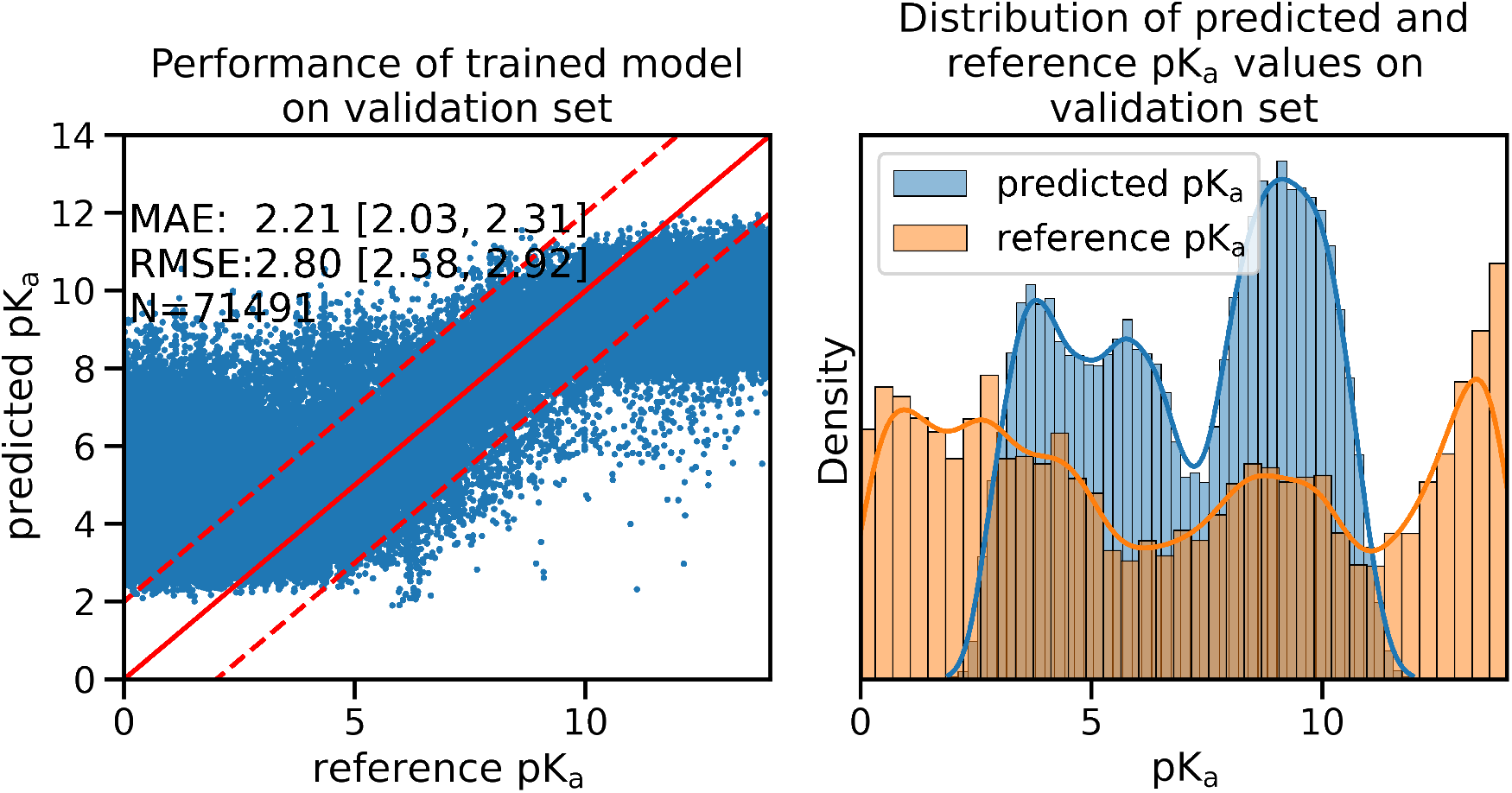
The performance of the fine-tuned GNN model on the ChEMBL data set is shown. In contrast to the results obtained with the fine-tuned models shown in Figure S.I.4 the models shown here did **not** use regularization. The performance of the GNN model decreased significantly on the ChEMBL data, shifting pK_*a*_ values above 12 and below 2. The solid red line in the scatter plot indicates the ideal behavior of the reference and predicted pK_*a*_ values, the dashed lines mark the ±1 pk_*a*_ unit interval. Mean absolute error (MAE) and root mean squared error (RMSE) are shown, the values in bracket indicate the 90% confidence interval calculated from 50 repetitions with random training/validation splits. *N* indicates the number of investigated samples.

**Figure S.I.7.**
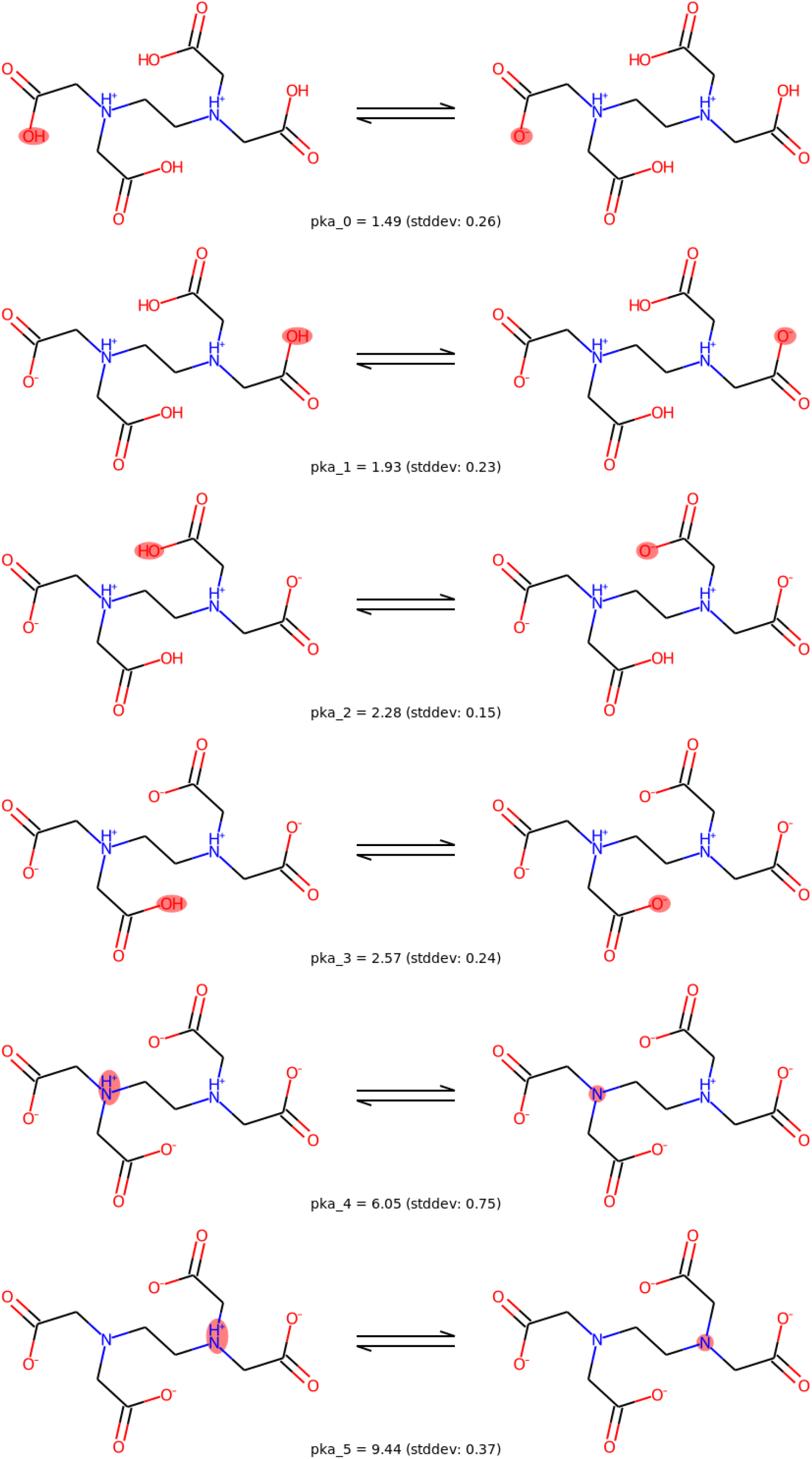
Results are shown for a sequential pK_*a*_ prediction using pkasolver for ethylenediaminetetraacetic acid (EDTA). For each protonation state the base-acid pair is shown and the consensus prediction for the pK_*a*_ value with the standard deviation is shown. The protonation site is highlighted for each protonation state.

## Footnotes

1 https://github.com/samplchallenges/SAMPL8

2 version 12.01, Advanced Chemistry Development Inc. 2010ACD/Labs

3 https://github.com/czodrowskilab/Machine-learning-meets-pKa

4 https://docs.scipy.org/doc/scipy/reference/generated/scipy.sparse.coo_matrix.html

